# Brainana: an end-to-end preprocessing framework for macaque neuroimaging

**DOI:** 10.64898/2026.06.03.729972

**Authors:** Xingyu Liu, Yijun Zhang, Zi Yin, Zonglei Zhen, Michael J. Arcaro

**Affiliations:** Department of Psychology, University of Pennsylvania, Philadelphia, PA 19104, USA; State Key Laboratory of Cognitive Neuroscience and Learning & IDG/McGovern Institute for Brain Research, Beijing Normal University, Beijing, 100875, China; Department of Psychological and Cognitive Sciences, Tsinghua University, Beijing, China; Beijing Key Laboratory of Applied Experimental Psychology, Faculty of Psychology, Beijing Normal University, Beijing, 100875, China

**Author notes:** Correspondence address: Xingyu Liu, Ph.D.

## Abstract

Macaque MRI bridges non-invasive systems neuroscience with cellular and circuit-level mechanisms, but preprocessing tools remain difficult to integrate and deploy reproducibly. We present Brainana, an automated, BIDS-compatible preprocessing and visualization framework for macaque neuroimaging. Brainana integrates structural and functional preprocessing, cortical surface reconstruction, quality control, transform tracking, and atlas projection within a containerized package, with cloud access for users without local compute. Macaque-trained deep learning models support brain extraction and tissue segmentation, while image orientation standardization and macaque-specific surface reconstruction optimizations address variability across acquisitions. A viewer automatically organizes derivatives and links volumetric and surface data, enabling users to inspect anatomy, cortical measures, atlas delineations, and activity maps without neuroimaging expertise. Across 23 imaging sites, Brainana processed heterogeneous data from 130 monkeys, yielding anatomical correspondence, reliable native-space surfaces, localized task-evoked activations, and reproducible brain-wide resting-state correlations. Brainana enables reproducible, scalable, and accessible macaque MRI analysis, cross-study comparison, and multimodal integration.

## Introduction

Macaque MRI provides an essential bridge between large-scale brain organization measured noninvasively in humans and the cellular and circuit-level mechanisms accessible only through invasive experiments ^1,2^. This connection supports mechanistic interpretation of human neuroimaging findings and enables direct tests of whether principles of brain organization generalize across species. Realizing this potential requires transforming raw MRI acquisitions into anatomically accurate, spatially consistent, and reproducible representations. Preprocessing is central to this transformation, correcting acquisition artifacts, aligning data across individuals, and defining the spatial and temporal structure on which downstream analyses depend.

Preprocessing choices directly shape the data that enter analysis and can affect statistical inference, anatomical localization, and measurements of brain structure and function ^3,4^. As neuroimaging workflows have grown more complex, requiring many interdependent operations and methodological choices, study-specific pipelines can introduce uncontrolled variability, limit cross-dataset comparability, and create barriers to reproducibility and reuse. In human neuroimaging, these concerns have motivated standardized, automated frameworks such as fMRIPrep ^5^ and DeepPrep ^6^, which integrate established tools into analysis-agnostic workflows that adapt to diverse datasets while improving transparency, reproducibility, and quality control.

Macaque neuroimaging lacks a comparable end-to-end framework. Important tools address individual components of macaque MRI analysis, including brain extraction, segmentation, registration, functional preprocessing, and cortical surface reconstruction ^7–12^, but these tools remain distributed across separate software packages. This fragmentation is compounded by challenges that prevent direct transfer of human pipelines to macaque data, including smaller brain size, species-specific morphology and tissue contrast, heterogeneous acquisition protocols, specialized coils, anesthesia, implanted hardware, and contrast agents ^9,10,12^. As a result, macaque-specific solutions have remained limited in scope, interoperability, or scalability.

The need for integrated preprocessing is especially acute in macaque studies that combine MRI with invasive experiments, the dominant use case for macaque neuroimaging. In many laboratories, MRI is acquired primarily to guide electrophysiology, pharmacological manipulation, stimulation, or lesion studies. These scans are essential for targeting and localization, but they could also provide the anatomical scaffold needed to link invasive measurements with atlas parcels, functional maps, and shared template spaces. Without accessible, end-to-end preprocessing, these data often remain confined to individual animals or study-specific coordinate systems, limiting quantitative analysis, cross-animal comparison, or integration across laboratories. Generating standardized derivatives, however, does not by itself make them straightforward to use. Researchers must still identify the appropriate files, load volumetric and surface data into specialized software, configure overlays and color maps, and relate corresponding locations across representations. These downstream technical demands can prevent non-specialists from inspecting processed data and applying them to anatomical localization, experimental planning, and scientific interpretation.

Here we introduce Brainana, a unified preprocessing framework for macaque MRI that brings core design principles from contemporary human neuroimaging pipelines to the non-human primate domain. Brainana integrates macaque-specific structural and functional preprocessing, cortical surface reconstruction, quality control reporting, transform tracking, and template-to-individual atlas projection within a single automated pipeline built around standardized data structures, reusable derivatives, and reproducible execution. Brainana is accompanied by an interactive viewer that automatically identifies pipeline outputs and provides linked visualization across volumetric and cortical surface representations, reducing the expertise required to inspect and use the resulting data. By combining macaque-specific methods with scalable workflow management, containerized deployment, and accessible visualization, Brainana provides an end-to-end platform for robust preprocessing across sites, acquisition protocols, and experimental contexts. In doing so, Brainana addresses a central infrastructure gap in macaque neuroimaging and provides a shared framework for linking MRI with physiology, intervention-based experiments, and multimodal data across spatial scales, from cellular measurements to brain-wide organization.

## Results

### An end-to-end pipeline for standardized and efficient macaque MRI preprocessing

Brainana processes structural MRI (sMRI) and functional MRI (fMRI) within a unified modular pipeline (Fig. 1A). The structural workflow uses T1-weighted (T1w) images as the primary anatomical reference, with optional T2-weighted (T2w) images, and performs volumetric preprocessing, brain extraction, tissue segmentation, spatial registration, and cortical surface reconstruction. The functional workflow processes blood oxygenation level dependent (BOLD) and monocrystalline iron oxide nanoparticles (MION) based T2*-weighted time series, performing run-level preprocessing, brain extraction, and registration either through subject-specific T1w space or directly to template space when subject-specific anatomical images are unavailable. This design enables consistent processing across datasets ranging from structural-only acquisitions to combined multimodal studies. Each stage can be run as part of the full pipeline or customized through user-defined parameters, allowing Brainana to maintain standardized processing while accommodating dataset-specific needs.

**Figure 1.**
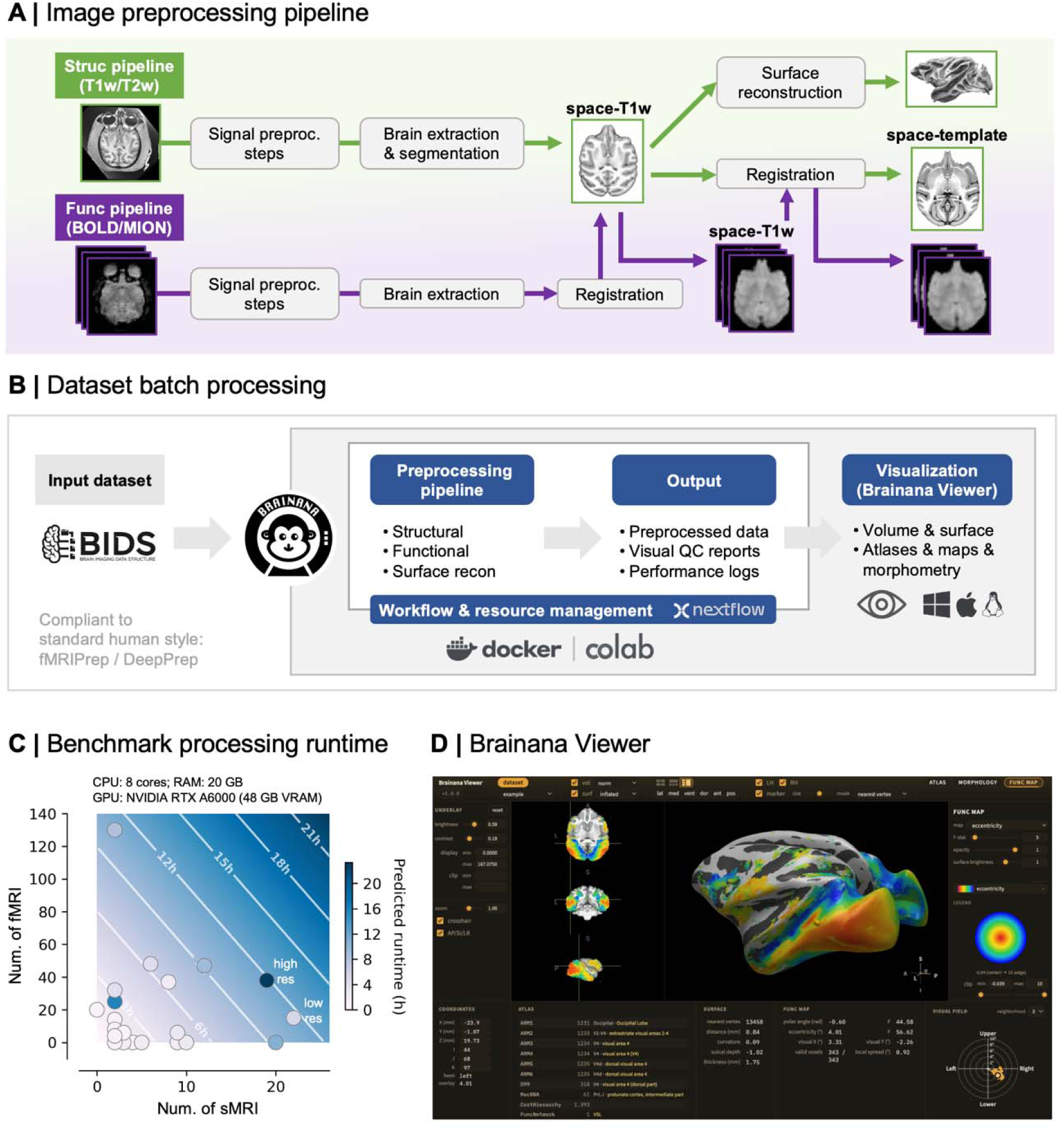
Overview, architecture, and performance of the Brainana pipeline. **(A) Brainana preprocessing pipeline.** Structural (top) and functional (bottom) workflows follow a shared sequence of image preprocessing, brain extraction, and spatial registration. In addition to currently available tools, the structural workflow performs tissue segmentation and cortical surface reconstruction using individual monkey cortical morphology. **(B) Brainana execution framework.** BIDS-formatted datasets are automatically parsed to identify jobs across subjects, sessions, and image types. Jobs are then routed to the modular workflows and executed in parallel with Nextflow^14^. Outputs include preprocessed images, spatial transforms, performance logs, and HTML quality control (QC) reports. **(C) Runtime performance across macaque MRI datasets.** Datasets from 20 sites spanning PRIME-DE, UNC-Wisconsin, and in-house collections were processed using the default full preprocessing configuration, including structural and functional volumetric preprocessing and surface reconstruction. Individual points represent sites, with structural image count on the x axis and functional image count on the y axis. Background shading and contour lines show predicted runtime in hours under the specified hardware setup. **(D) Brainana Viewer interface**. The viewer automatically displays Brainana derivatives in linked volumetric and surface views, including cortical morphometric measures, atlas labels, and template-derived functional maps projected into individual space. Here, a template-projected eccentricity map is displayed on an indivudal monkey’s anatomy together with the visual field coverage associated with the selected cortical location.

Brainana follows the organizational principles of contemporary human MRI pipelines, such as fMRIPrep ^5^ and DeepPrep ^6^, while implementing a macaque-specific preprocessing framework (Fig. 1B). Datasets formatted according to the Brain Imaging Data Structure (BIDS)^13^ are automatically parsed to identify subjects, sessions, modalities, and metadata, eliminating manual specification. An initialization step generates structured preprocessing jobs, which are then routed through modular structural, functional, and surface reconstruction workflows. The workflow manager Nextflow orchestrates parallel execution, dependency tracking, caching, and restart behavior across local workstations, clusters, and other compute environments. Brainana is distributed as a Docker container with fixed software versions, eliminating the need for users to install and configure individual dependencies and ensuring a consistent runtime environment across machines. For users without local GPU resources, Brainana Lite provides a cloud-accessible version of the volumetric structural workflow through Google Colab, enabling GPU-accelerated structural preprocessing overcoming local hardware constraints.

The pipeline generates standardized derivatives, including preprocessed images, spatial transforms, segmentation maps, cortical surfaces, performance logs, and structured HTML quality control (QC) reports. These outputs are designed to support both routine preprocessing and downstream reuse of spatial mappings, atlases, and QC information. Across 23 macaque MRI sites, a representative subject with one sMRI and one fMRI required approximately 30 minutes to process on a modern local workstation, with GPU-enabled runs completing up to approximately 60% faster depending on dataset composition (Fig. 1C; Supp. Fig. 1). Together, Brainana provides a scalable and computationally efficient framework for applications ranging from single-subject analyses to large multi-site macaque MRI datasets.

Brainana further provides an integrated visualization app, Viewer (Fig. 1D), that makes derivatives accessible for downstream use without requiring familiarity with neuroimaging file formats, naming conventions, or dedicated visualization software. Viewer automatically identifies and loads the relevant outputs for each subject, presenting anatomical volumes, cortical surfaces, and surface-derived measures such as thickness and curvature without manual file selection or configuration. Volumetric and surface views are spatially linked, allowing users to relate corresponding locations across representations, and individual data can be viewed alongside atlases and reference maps projected from template space, including group-derived visual-field coverage maps back-projected into subject anatomy. By automating data discovery, display configuration, and cross-representation correspondence, Viewer lowers the barrier between generating neuroimaging derivatives and using them for anatomical localization, experimental planning, and scientific interpretation. It is distributed as a native app for macOS, Windows, and Linux.

### Structural preprocessing resolves macaque-specific acquisition variability

The structural preprocessing workflow handles variability typical of macaque MRI across laboratories, including sources of heterogeneity that commonly cause standard MRI analysis tools to fail, and produces accurate sMRI in native and template spaces, together with brain masks and tissue segmentations (Fig. 2A). When multiple T1w images are available within a session, they are co-registered and averaged into a single reference volume. For longitudinal datasets, sessions can be processed separately to preserve session-specific anatomy or aligned to a reference session. Optional T2w images are aligned to this reference and carried through subsequent processing steps.

**Figure 2.**
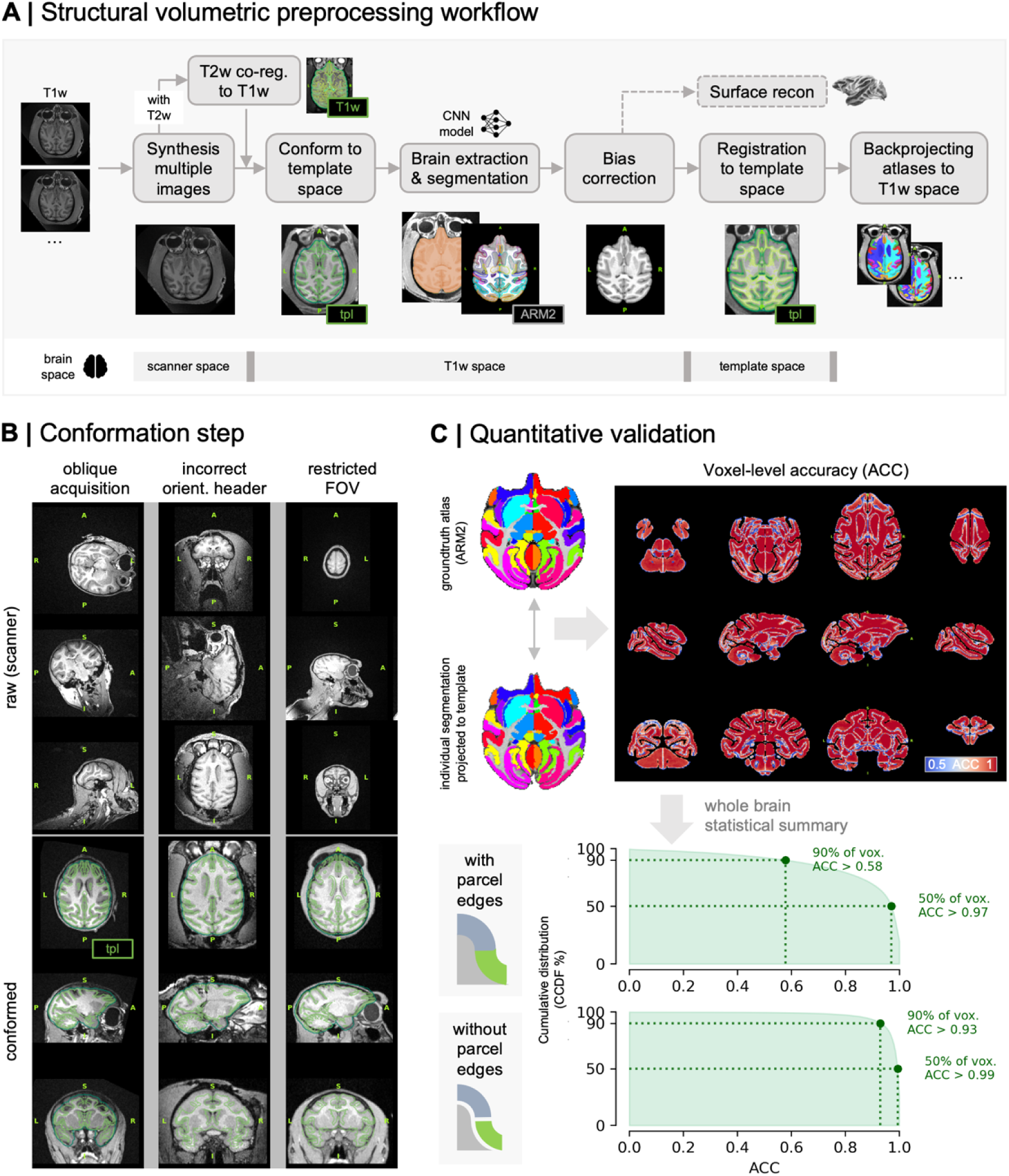
Structural preprocessing and validation. **(A) Structural volumetric preprocessing workflow.** Multiple T1w scans, when available, are combined into a reference volume. Optional T2w images are co-registered to the T1w reference. Processing then proceeds through conformation to template orientation, CNN-based brain extraction and ARM2 segmentation, bias field correction, and nonlinear registration to template space. Template-space atlases are back-projected to individual T1w space using the estimated transforms. Cortical surface reconstruction uses the preprocessed T1w image and segmentation outputs. Contours from reference images, such as the T1w image, template, or atlas, are overlaid on the corresponding data to facilitate visual QC. **(B) Conformation of challenging macaque MRI acquisitions.** Example T1w images show common acquisition issues (top), including oblique orientation, incorrect orientation metadata, and restricted field of view caused by head-post hardware or variable subject positioning, alongside the corresponding conformed images in standard orientation (bottom). The conformation step brings images into standard orientation and approximate template position using rigid registration without altering brain shape. Contours in conformed images indicate the template. **(C) Quantitative validation of tissue segmentation.** Individual ARM2 segmentations were propagated to template space and compared against the template-space ARM2 atlas. The voxel-level accuracy map shows the proportion of monkeys with correctly labeled voxels on the NMT2Sym template. Accuracy is high throughout the brain, with lower values confined to parcel boundaries.

A key feature of the structural workflow is a conformation step that resolves common inconsistencies in macaque MRI data (Fig. 2B). Unlike human MRI, macaque acquisitions frequently exhibit oblique orientations, incorrect header metadata, and restricted fields of view due to head-post hardware or variable animal positioning (Fig. 2B). These issues cause downstream tools that assume standard orientation to fail or require manual correction. Conformation avoids these failures by applying a rigid transform that places images in a standard orientation (RAS+) and approximate template position, while preserving the shape of the individual brain.

Brainana incorporates deep learning-based methods for brain extraction, tissue segmentation, and anatomical parcellation. The conformed T1w image is processed with a CNN adapted from FastSurferCNN ^15^, a model originally trained to segment human brains, which we then fine-tuned for macaque neuroanatomy. The model produces brain masks, hemisphere labels, and ARM2 anatomical labels, a composite parcellation derived from level 2 of the Cortical Hierarchy Atlas of the Rhesus Macaque (CHARM)^9^ and the Subcortical Atlas of the Rhesus Macaque (SARM)^16^. Brain-extracted T1w images are bias-corrected and nonlinearly registered to template space, with transforms retained for projecting template-space atlases and maps into individual subject space. The resulting transforms are used to project template-space atlases and maps into both T1w space and scanner space, linking individual anatomy to shared reference systems and enabling atlas-guided localization in the original acquisition coordinate frame.

We evaluated segmentation and registration accuracy by comparing subject-specific parcellations, propagated to template space, against the ARM2 template atlas (Fig. 2C). Voxel-level accuracy was defined as the proportion of correctly labeled voxels across subjects (N=97). Accuracy was high throughout the brain, with the strongest agreement in parcel interiors and expected reductions at parcel boundaries, where anatomical variability, partial volume effects, and registration uncertainty are greatest. After excluding boundary voxels, 90% of voxels exceeded an accuracy of 0.93, and 50% exceeded an accuracy of 0.99, indicating consistent voxel-level correspondence between subject-derived and template-space anatomical labels.

### Native-space cortical surface reconstruction enables morphometric analysis

Brainana uses preprocessed T1w images and segmentation outputs to reconstruct cortical surfaces in individual anatomical space, enabling analyses of cortical morphology (Fig. 3A). Surface reconstruction is performed using a workflow based on FastSurfer ^15^ and FreeSurfer ^17^, with four macaque-specific adaptations: (1) conversion of CNN-derived ARM2 labels into a FreeSurfer-style aparc+aseg segmentation for cortical reconstruction; (2) surface reconstruction parameters adjusted for submillimeter macaque MRI; (3) targeted refinements in regions prone to segmentation errors, including the occipital calcarine cortex, claustrum, and orbitofrontal cortex (Supp. Fig. 2); and (4) an additional topology correction step for surface defects that commonly arise in macaque reconstructions and are not reliably resolved by FreeSurfer alone. The targeted refinements improve segmentation quality in areas where thin white matter, low contrast, or anatomical complexity can disrupt accurate delineation of the grey-white boundary and pial surface, thereby reducing downstream surface reconstruction failures. Resulting cortical surfaces are also provided in GIFTI format to support platform-independent exchange and analysis.

**Figure 3.**
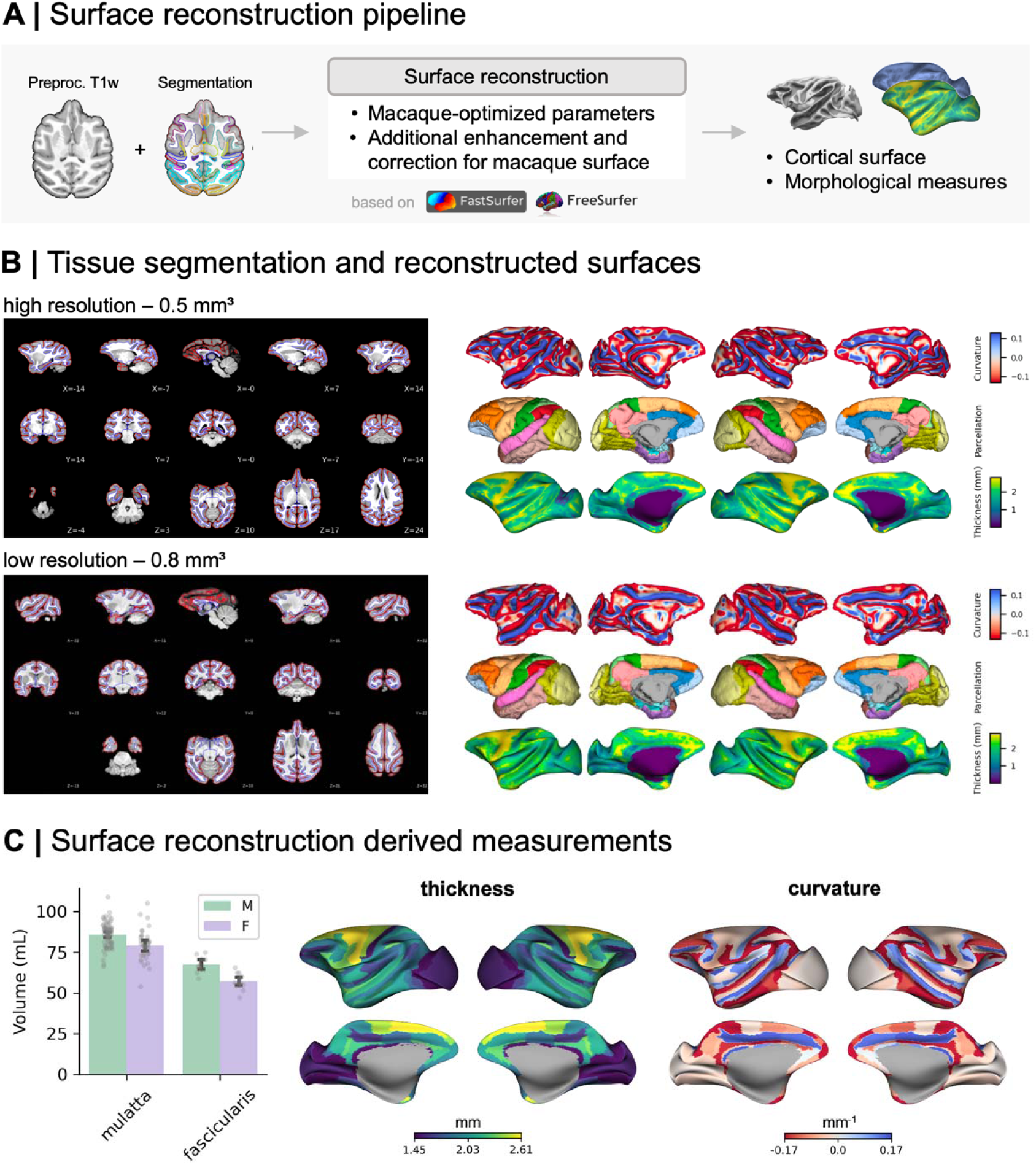
Cortical surface reconstruction and group-level morphology. **(A) Cortical surface reconstruction workflow.** Preprocessed T1w images and macaque-specific FreeSurfer-style aparc+aseg segmentations derived from ARM2 labels are used as inputs. Targeted refinements are applied before reconstruction to improve segmentation in regions that are challenging in macaque MRI, including regions with thin or low-contrast white matter, such as calcarine, claustrum, and orbitofrontal cortex. The refined inputs are then passed to a FastSurfer- and FreeSurfer-based reconstruction workflow with macaque-specific adaptations, producing cortical surfaces and associated morphological measures in individual native space. **(B) Example cortical surface reconstructions across image resolutions.** White matter (inner) and pial (outer) surfaces are shown in lateral, coronal, and axial views for macaque T1w images acquired at high resolution (0.5 mm³) and at lower resolution (0.8 mm³). Reconstructions are reliable across resolutions, enabling estimation of cortical curvature, thickness, and surface area across datasets. **(C) Group-level morphological outputs.** Measurements were derived from preprocessed T1w images across multiple datasets and sites. Total brain volumes are reported separately for Macaca mulatta and Macaca fascicularis groups. Average cortical thickness and curvature are projected onto the NMT2Sym template at CHARM level 6 for Macaca mulatta.

Surface reconstructions accurately delineate cortical boundaries across a range of image resolutions (Fig. 3B). White matter and pial surfaces can be reconstructed at both high-resolution (0.5 mm³) and lower-resolution (0.8 mm³) acquisitions, spanning the range commonly encountered in macaque MRI datasets. These surfaces enable the measurement of cortical curvature, thickness, and surface area in individual native space. At the group level, morphological measures derived from datasets acquired across 19 sites were consistent with known anatomical patterns (Fig. 3C). Total brain volume was 84 ± 9 mL in rhesus macaques (N = 107) and 62 ± 7 mL in cynomolgus macaques (N = 23), consistent with reported species differences in brain size ^18–20^. Cortical thickness maps showed thicker cortex in frontal regions and thinner cortex in occipital regions, consistent with prior work ^21,22^, and curvature maps delineated major gyral and sulcal anatomy (Fig. 3C). Together, these results demonstrate that Brainana enables accurate and consistent surface reconstruction for individual- and group-level morphological analyses.

### Functional preprocessing preserves localized task-evoked responses and brain-wide resting-state organization

The functional preprocessing workflow standardizes analysis of BOLD and MION-based time series while adapting the registration strategy to the available structural reference (Fig. 4A; Supp. Fig. 3). Each functional run undergoes slice timing correction followed by motion correction. For sessions with multiple runs, all runs are co-registered to a designated reference run to ensure within-session spatial correspondence in scanner space. A session-level representative image is generated by averaging across time points and runs, and then conformed to the selected structural reference using the same procedure as the structural workflow. Brain extraction is applied to the conformed representative image using a CNN model. When a subject-specific T1w image is available, the functional reference is registered to T1w space and projected to template space using transforms from the structural workflow. When no T1w image is available, the functional reference is registered directly to template space (Supp. Fig. 3).

**Figure 4.**
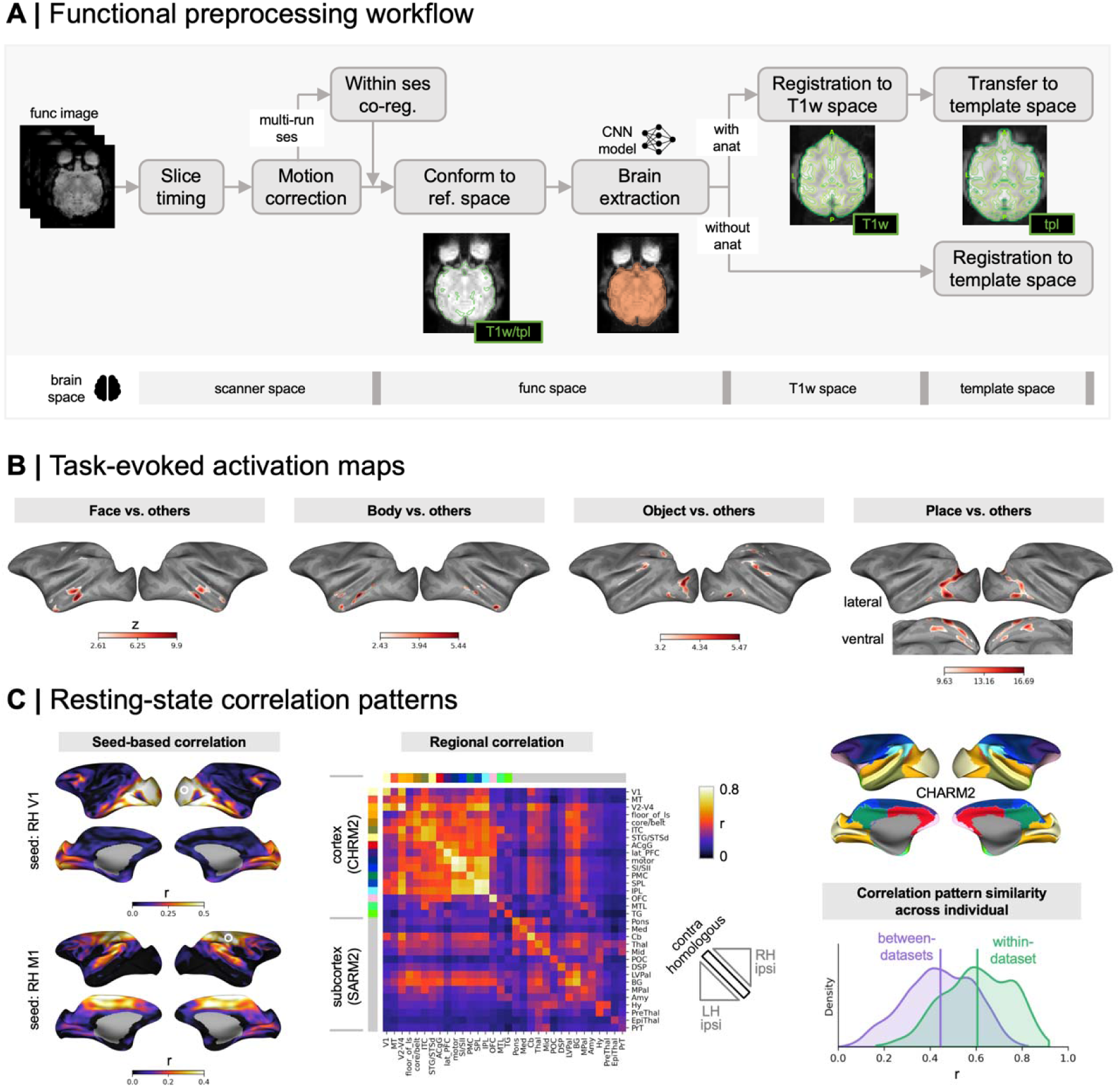
Functional preprocessing and validation of functional organization. **(A) Functional preprocessing workflow.** Each functional run undergoes slice timing correction and motion correction. For multi-run sessions, runs are co-registered to a reference run to ensure within-session alignment in scanner space. A session-level representative FUNC image is derived, conformed to the selected anatomical reference, and brain extracted using a CNN model. If a subject-specific T1w image is available, the FUNC reference is registered to T1w space and projected to template space using T1w-to-template transforms; otherwise, the FUNC reference is registered directly to the template. **(B) Task-evoked category-selective activation maps from a visual localizer experiment.** Statistical maps show contrasts comparing each visual category with all other categories for faces, bodies, objects, and places. Contrast maps are shown for an example macaque and displayed at the top 5% of vertices. Responses appeared in expected category-selective regions of macaque visual cortex, supporting accurate functional preprocessing and registration. **(C) Resting-state correlation validation across individuals and datasets.** (Left) Seed-based maps from a single representative macaque, showing correlations across the cortical surface for seeds in primary visual cortex (V1) and primary motor cortex (M1). (Middle) Group-level regional matrix computed across ARM2 regions, showing stronger correlations between contralateral homologous regions, consistent with expected hemispheric symmetry. (Right) Similarity of regional correlation patterns between pairs of macaques. Each point reflects the similarity between two monkeys’ region × region correlation matrices, comparing pairs drawn from the same dataset and from different datasets. Similarities are positive in both cases, indicating reproducible resting-state correlation structure across individuals, and are higher within datasets than between datasets, consistent with additional sensitivity to dataset-specific acquisition and cohort factors.

To validate the functional preprocessing workflow, we used two complementary datasets that probe different requirements for accurate fMRI preprocessing. A visual localizer dataset ^23^ tested whether Brainana preserves spatially precise, localized task-evoked activations after motion correction, brain extraction, functional-to-anatomical alignment, and template registration. A larger resting-state dataset from 78 macaques across 13 sites ^2^ tested whether Brainana preserves brain-wide correlation structure across individuals, datasets, and acquisition protocols. In the visual localizer, contrasts comparing each category with all other categories revealed face-, body-, object-, and place-selective responses in the expected regions of macaque visual cortex (Fig. 4B)^23–27^, illustrating accurate functional-to-anatomical alignment and template registration.

Seed-based correlation maps for primary visual cortex (V1) and primary motor cortex (M1) in an individual macaque showed distinct cortical patterns consistent with their different functional networks (Fig. 4C). At the group level, regional correlation matrices showed elevated correlations among homologous regions across hemispheres (Fig. 4C), consistent with expected hemispheric symmetry. Correlation structure was reproducible across individuals, with positive similarity for both within-dataset and between-dataset comparisons. Similarity was higher for pairs from the same dataset than for pairs from different datasets, consistent with additional sensitivity to acquisition parameters, image properties, or cohort differences. Together, these results show that the functional preprocessing workflow preserves both spatially localized task-evoked activations and brain-wide resting-state organization.

### Integrated templates, atlases, and brain maps support cross-study integration

Brainana provides an integrated collection of templates, atlases, and brain maps that supports accessibility, reproducibility, and cross-study comparison in macaque neuroimaging (Fig. 5A). The collection includes five macaque anatomical templates: NMT v2 Sym, NMT v2 Asym ^9,28^, MEBRAINS ^29^, Yerkes19 ^30^, and D99 ^31^, each paired with volumetric and surface representations, allowing analyses to be performed consistently across different reference spaces. Anatomical and functional atlases include the ARM parcellation across six hierarchical levels (ARM1 to ARM6) ^9,16^, the MacBNA atlas ^32^, and functional network parcellations ^33,34^ (Fig. 5B). Additional brain maps capture retinotopic organization ^35^, somatotopic organization ^36^, functional connectivity gradients ^34^, as well as other features of macaque brain organization.

**Figure 5.**
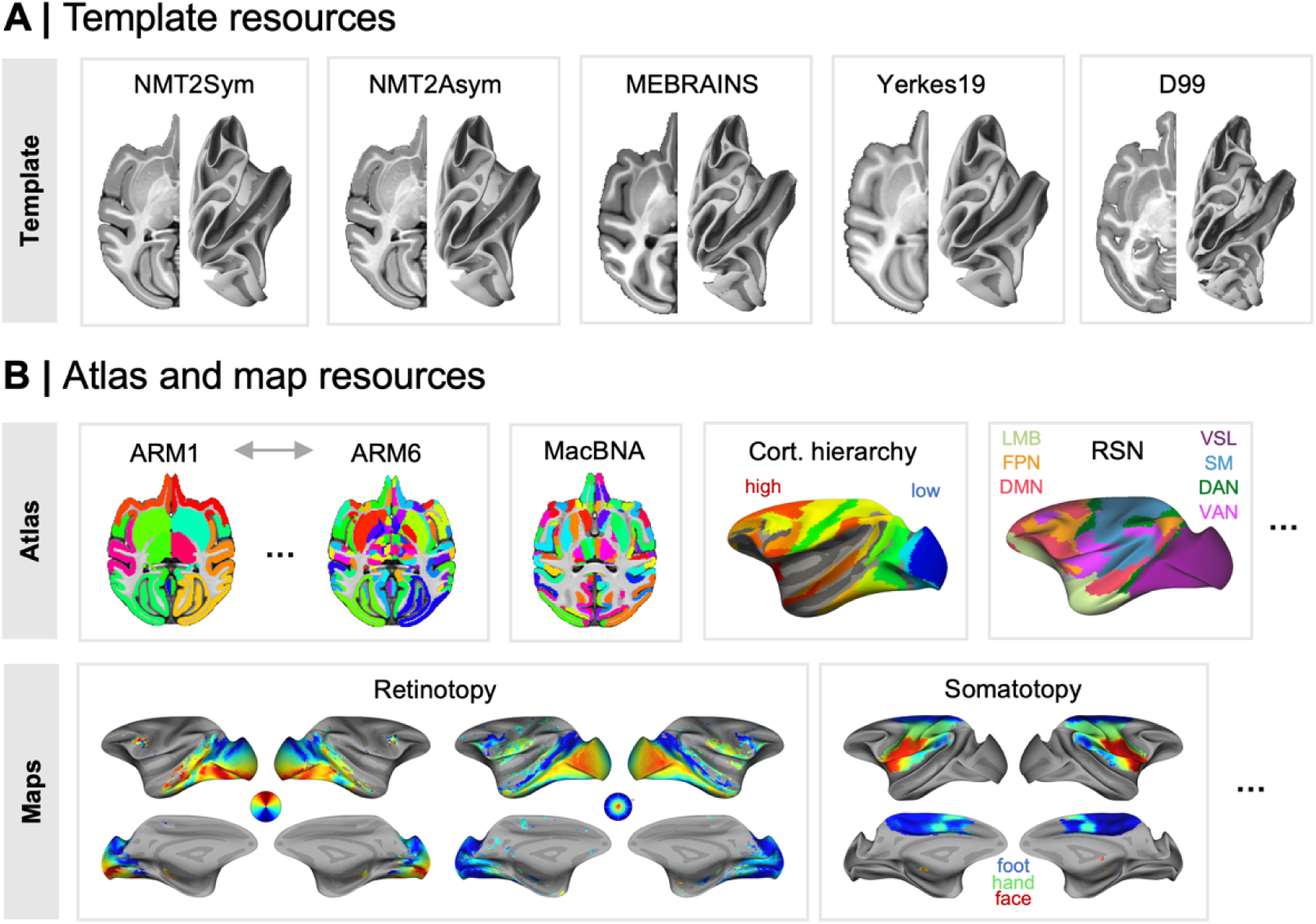
Integrated template, atlas, and brain map resources. **(A) Brainana template resources.** Brainana includes five macaque anatomical templates in volumetric and surface spaces: NMT v2 Sym, NMT v2 Asym, MEBRAINS, Yerkes19, and D99. **(B) Brainana atlas and map resources.** The collection encompasses anatomical and functional atlases and maps, including hierarchical parcellations (ARM1–ARM6), functional network parcellations, and maps of retinotopic organization, somatotopic organization, and other features of macaque brain organization.

These resources are directly integrated with Brainana’s preprocessing workflows, allowing template-space atlases and maps to be projected into individual T1w and scanner space using the estimated spatial transforms, without requiring separate registration or atlas-projection steps. Because transforms are explicitly tracked and exported, anatomical regions, functional maps, and experimental measurements can be related within a common coordinate system, ensuring consistency across data types and supporting quantitative comparison across animals, sessions, and laboratories.

### A unified framework for the broader macaque neuroscience community

Brainana is designed to promote integration of macaque MRI with physiology, histology, and intervention-based experiments, extending its utility beyond preprocessing (Fig. 6A). Many laboratories acquire MRI to guide recordings, stimulation, or lesion placement, but these scans are often used only for the original experiment. Brainana extends the value of these data by placing each animal’s MRI in both individual and template spaces and preserving the transforms needed to map between them. This provides a common anatomical frame for relating experimental measurements to individual anatomy, atlas parcels, and group-level reference maps. For example, CT images can be co-registered to the preprocessed T1w image to localize electrodes relative to the animal’s anatomy and atlas parcels projected into individual space, allowing recording sites to be described in anatomical terms rather than only by chamber position or electrode depth. This makes physiological data easier to interpret, share, and compare across animals and laboratories.

**Figure 6.**
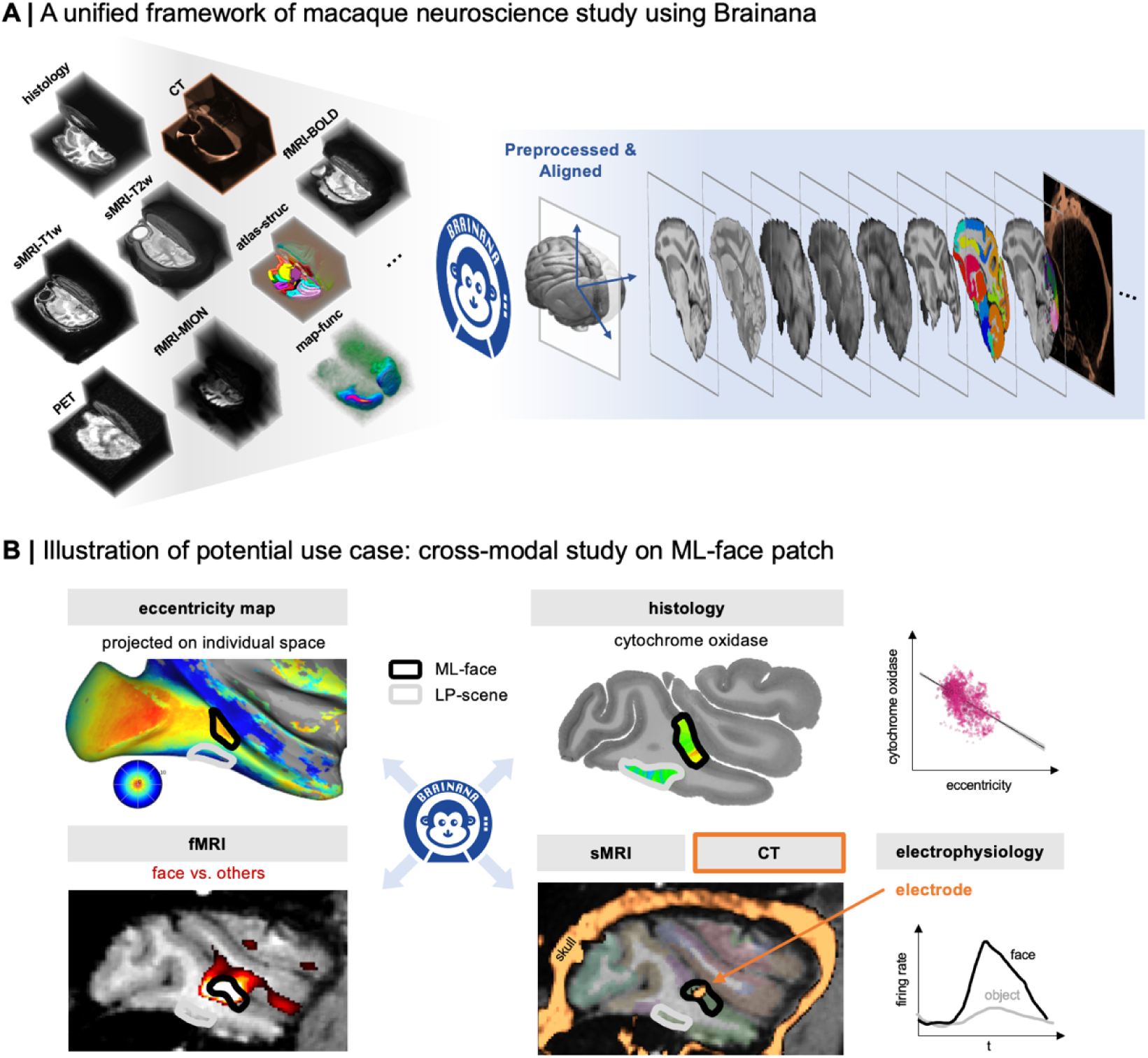
Brainana as a unified framework for the broader macaque neuroscience community. **(A) A unified framework for macaque neuroscience study using Brainana.** Data from diverse modalities, including sMRI, fMRI, CT, PET, histology, atlases, and functional maps, can be linked through standardized individual and template spaces, extending Brainana’s utility beyond MRI preprocessing and enabling cross-modal analyses of the macaque brain. **(B) An illustration of multi-scale, cross-modal integration centered on the middle lateral face patch (ML-face).** Adapted and assembled from published data. Co-registering retinotopic maps ^35^, face-selective fMRI activation, cytochrome oxidase histology ^37^, and CT-localized electrodes^38^ into a shared individual space demonstrates how Brainana can connect macroscale functional imaging, metabolic architecture, and single-unit electrophysiology within a shared anatomical reference frame.

A multimodal characterization of the middle lateral face patch (ML-face), assembled from previously published data, illustrates how Brainana supports cross-modal integration across spatial scales, from macroscale brain architecture to cellular signaling (Fig. 6B). Co-registering retinotopic maps ^35^, face-selective fMRI activation, cytochrome oxidase histology ^37^, and CT-localized electrodes ^38^ into a shared individual space provides a common anatomical reference for linking these modalities. Within this reference, fMRI activation localizes the face-selective ML region, retinotopic maps characterize the spatial coding of receptive fields within it, CT-localized electrodes confirm spatial and feature selectivity at single-unit and population level, and cytochrome oxidase histology reveals its metabolic architecture. Together, these linked measurements support experimental planning, post-hoc localization, and comparison across approaches for studying brain organization.

## Discussion

Brainana demonstrates that standardized, end-to-end preprocessing can be implemented for macaque MRI despite the acquisition heterogeneity that has historically required bespoke, study-specific workflows. The BIDS-compatible pipeline coordinates structural, functional, and surface reconstruction workflows with atlas projection, quality control, and explicit transform tracking, producing linked outputs across individual, template, and surface spaces. Across datasets from 23 imaging sites spanning two macaque species and diverse acquisition protocols, Brainana recovered expected anatomical structure, preserved spatially localized task-evoked visual responses, and maintained reproducible brain-wide resting-state correlation patterns.

Existing tools have advanced individual components of macaque MRI analysis, including segmentation, registration, functional preprocessing, and surface reconstruction ^7–12^, but these capabilities often remain distributed across separate software packages and study-specific workflows. Brainana addresses this fragmentation by bringing these capabilities into a single BIDS-compatible pipeline while adding macaque-trained deep learning models, a conformation procedure for resolving acquisition inconsistencies common in macaque imaging, and surface reconstruction refinements for regions that are challenging to segment in macaque brains. By maintaining outputs in individual anatomical space and template space, the pipeline supports subject-specific analyses, cross-subject comparison, and consistent use of common anatomical and functional reference systems.

Brainana addresses a practical barrier that has limited the use of macaque MRI beyond its immediate experimental purpose. Many laboratories acquire MRI to guide localization in electrophysiology, stimulation, pharmacological manipulation, or lesion experiments, but these scans are rarely processed further because doing so requires specialized software expertise, dependency management, and neuroanatomical knowledge. Brainana makes this additional processing practical by automating the generation of preprocessed structural images, spatial transforms, and atlas projections. These outputs allow experimental measurements to be related to individual anatomy, template coordinates, atlas labels, and functional maps, supporting quantitative analysis and comparison across sessions, animals, laboratories, and studies.

Brainana Viewer extends this accessibility beyond preprocessing by automatically organizing these derivatives and displaying linked volumetric and surface representations together with morphometric measures, atlases, and reference maps. This integration reduces the specialized knowledge otherwise needed to select derivative files, configure visualization software, and establish correspondence across coordinate systems. The combined framework therefore supports researchers who need neuroimaging for localization, experimental planning, or interpretation without requiring them to become expert users of multiple neuroimaging software packages. Brainana Lite further lowers the barrier to adoption by enabling cloud-based structural preprocessing for users without local infrastructure.

Several directions remain to extend Brainana’s scope, generalizability, and community utility. First, deeper integration of additional modalities would expand support for experimental targeting, molecular imaging, and cross-modal analyses beyond the core cases demonstrated here (Fig. 6). Second, current structural and surface workflows are optimized for adult macaque T1w contrast. Neonatal and early developmental datasets, where tissue contrast differs substantially from adults, will require additional training data and model adaptation. Third, the framework is designed to be extensible to other NHP species, but species-specific training and validation will be needed. Finally, a species-representative group surface template analogous to fsaverage ^39^ or onavg ^40^ would further standardize surface-based group analyses and complement the individual-space surfaces Brainana currently produces. Continued development alongside community resources such as PRIME-RE ^1^ and TemplateFlow ^41^ will be important for long-term accessibility, interoperability, and adoption.

Realizing the translational potential of macaque neuroimaging requires preprocessing infrastructure that can accommodate the field’s diversity of datasets, acquisition protocols, and experimental contexts. Through an open, containerized preprocessing workflow and integrated visualization app, Brainana links standardized MRI derivatives with anatomical reference systems, functional maps, and multimodal experimental measurements. By making these linked outputs easier to generate, inspect, and interpret, this design enables integrative analyses across animals, laboratories, modalities, and spatial scales, with the flexibility to grow alongside community needs and the expanding role of macaque MRI in neuroscience.

## Online Methods

### Datasets for model and pipeline evaluation

Three datasets were used to train and evaluate Brainana. Detailed acquisition and demographic information are provided in the original publications; brief summaries are provided below.

#### PRIME-DE

The PRIME-DE open-resource dataset ^2^ aggregates macaque MRI data from 23 sites and includes both *Macaca mulatta* and *Macaca fascicularis* across a broad age range.

Sites vary in available modalities, contributing sMRI, fMRI, or both. Acquisition parameters vary substantially across sites in scanner vendor, model, and field strength, as well as sequence-level differences in spatial resolution, field of view, orientation and image contrast. This heterogeneity was used to evaluate Brainana across the range of data encountered in the macaque MRI community.

#### In-house dataset

An in-house dataset ^35,38,42^ comprised 17 animals scanned on a 3 T Siemens Tim Trio scanner. T1w images were acquired using an MP-RAGE sequence with TR = 2.6 s, TE = 4.9 ms, and 0.5 mm isotropic resolution. fMRI was acquired with TR = 2 s, TE = 13 ms, and 1 mm isotropic resolution following injection of the MION contrast agent. For evaluation of task-evoked responses, 11 visual localizer runs collected from a single session of one animal were analyzed.

#### UNC-Wisconsin dataset

The UNC-Wisconsin Rhesus Macaque Neurodevelopment Database ^43^ provides sMRI data from 34 typically developing macaques scanned longitudinally from 2 weeks to 36 months of age. T1w images were acquired on a GE MR750 3.0 T scanner using a fast gradient echo sequence (GE BRAVO; TR = 8.684 ms, TE = 3.652 ms, effective resolution 0.55 x 0.55 x 0.80 mm). T2w images were acquired using a 3D CUBE FSE sequence (TR = 2500 ms, TE = 87 ms, effective resolution 0.60 x 0.60 x 0.60 mm).

### Configuration and workflow dispatch

Brainana is controlled by a YAML configuration file that specifies dataset filters, target template space, enabled processing branches, algorithm-specific parameters, and registration settings. Users can restrict processing to selected monkeys, sessions, tasks, or runs; select the output template space; enable or disable structural processing steps (synthesis, conformation, segmentation, and bias correction), surface reconstruction, and functional processing steps (slice timing correction, motion correction, functional bias correction, and brain extraction); and specify registration types and interpolation.

Before execution, Brainana scans the input BIDS directory and generates structured job descriptors for each monkey, session, modality, and image type. These descriptors specify input paths, BIDS entities, available metadata, and required processing branches. The job descriptors are passed to Nextflow as channels that instantiate only the relevant workflow components for each dataset.

### T1w brain extraction and segmentation model

#### Model training

A dataset of 121 T1w images was assembled from PRIME-DE, the in-house dataset, and UNC-Wisconsin. To prevent individual sites from dominating training, per-site contributions were capped at 18 images. Images were split 80% training and 20% validation sets.

Training labels were generated by warping the ARM2 atlas from template space to individual subject space. Specifically, initial brain extraction was performed with DeepBet ^44^, followed by nonlinear registration to the NMT2Sym template using ANTs. Registered images and propagated labels were visually inspected, and only cases passing quality control inspection were retained for model training.

We adapted the human brain-trained FastSurferCNN ^15^ for macaque T1w segmentation by training plane-specific 2D models in the ARM2 label space. The axial and coronal models used 71 classes, and the sagittal model used 36 classes collapsing contralateral homologues into single labels. Training-time augmentation included random scaling, translation, additive Gaussian noise, and simulated MRI bias fields. The loss function combined softmax Dice loss and spatially weighted cross-entropy with equal weights. Models were trained using AdamW optimization with cosine annealing. The mean validation Dice score across planes was 0.85.

#### Model inference within structural workflow

Trained models are packaged and deployed within the structural workflow to perform segmentation and brain extraction during execution. For each input T1w image, plane-specific predictions are generated independently and combined using weighted fusion, with weights of 0.4 for coronal, 0.4 for axial and 0.2 for sagittal outputs.

The fused segmentation produces ARM2 labels, from which whole-brain and hemisphere masks are derived using morphological post-processing that fill in holes through dilation and erosion.

## Structural preprocessing

The structural workflow produces bias-corrected, brain-extracted, segmented, and template-registered outputs from raw T1w images, with optional T2w support. When multiple T1w (or T2w) images are available, Brainana synthesizes a single anatomical reference by rigidly co-registering all images to the lexicographically first image and averaging in reference space.

The T1w image in scanner space is conformed to the selected template orientation and grid to better enable subsequent segmentation and registration. Specifically, the input is first brain-extracted with DeepBet ^44^, then registered to the template via FSL FLIRT (6 DOF rigid), and the resulting transform is applied to the full-head image. This produces the conformed T1w image (T1w space) and writes transforms between scanner and T1w space.

The conformed T1w image is segmented using the macaque-adapted FastSurferCNN model described above. The segmentation provides ARM2 labels, a brain mask, and hemisphere masks. The brain mask is applied to generate a brain-extracted T1w image. Bias field correction is then performed with ANTs N4BiasFieldCorrection and is restricted to brain tissue using the brain mask.

The T1w brain image is then registered to template space using a configurable multi-stage ANTs registration procedure. Registration proceeds through translation, rigid, affine, and optional SyN stages. When GPU resources are available, the SyN stage can be accelerated with FireANTs. Forward and inverse transforms are retained for subsequent atlas projection and functional registration.

When T2w images are available, they are rigidly co-registered to the preprocessed T1w image and resampled into T1w space. T2w images can then be carried into template space using the T1w-derived transforms.

### Surface reconstruction

Brainana adapts the FastSurfer ^15^ framework for macaque cortical surface reconstruction using a two-stage workflow. First, a macaque-adapted FastSurferCNN described above generates volumetric T1w segmentations. These segmentations are then used to initialize cortical surface reconstruction and morphometry. This approach preserves FreeSurfer-compatible ^39^ inputs and outputs while replacing human-specific atlas priors with macaque-specific ARM2 labels and additional species-specific refinements.

Before surface reconstruction, Brainana applies targeted refinements to improve segmentation in regions that are prone to reconstruction errors in macaque MRI. First, medial pallium and amygdala labels are reclassified as cortex to support accurate delineation of anterior temporal and parahippocampal regions. Second, white matter segmentation is refined in the occipital calcarine sulcus, where thin white matter often appears spatially discontinuous, and in orbitofrontal cortex, where local contrast and midline anatomy can produce missing white matter labels. For the calcarine sulcus, a more liberal white matter mask is derived from the CHARM segmentation, dilated to recover missing tissue, projected from template to individual space, and incorporated into the individual segmentation. In the orbitofrontal cortex, an analogous correction is applied without dilation, as CHARM-based labels alone are sufficient to recover missing white matter.

#### Surface reconstruction and morphometry

The refined T1w image and segmentation are passed to a FreeSurfer-style staged workflow covering volume preparation, bias correction, mask and segmentation preparation, white matter filling, tessellation, smoothing, inflation, spherical projection, topology correction, parcellation mapping, white and pial surface placement, cortical ribbon generation, and morphometric output. Four macaque-specific modifications were implemented:

1. The claustrum and immediately surrounding subcortical structures are masked and their voxel intensities are raised to the local white matter mean +10. This prevents the surface from blending into subcortical tissue, a common problem because the macaque claustrum is thin and low contrast in T1w images.
2. Submillimeter voxel spacing, typical of macaque MRI, is detected automatically and triggers high-resolution FreeSurfer processing to preserve anatomical detail.
3. A progressive mesh relaxation strategy is applied to the face-area remeshing step, relaxing constraints only when default thresholds fail. This avoids mesh instability arising from the relatively small cortical extents of macaque hemispheres while preserving geometric detail.
4. An additional iterative mesh repair step is applied when FreeSurfer’s built-in topology correction does not fully resolve surface defects, continuing until a valid manifold surface is obtained.

Surface outputs are written in both FreeSurfer-compatible and GIFTI formats. The GIFTI files provide standardized, platform-independent surface representations for downstream visualization, exchange, and analysis. During conversion, surface coordinates are corrected so the surface meshes align with the volume data.

### Functional brain extraction model

#### Model training

A dataset of 88 fMRI images was assembled from PRIME-DE, the in-house dataset, and UNC-Wisconsin, with per-site contributions capped at 10 images. Images were split into 80% training and 20% validation sets.

Training labels were generated in two stages. A precursor model was first fine-tuned using six manually labeled brain masks of functional data and then applied to the remaining functional images to generate approximate masks. Each coarsely brain-extracted functional image was rigidly registered to its corresponding T1w image, and the T1w brain mask was projected back to functional space to serve as the training label. Registered images and masks were manually inspected, and only cases passing QC were retained.

The functional brain extraction CNN was adapted from DeepBet ^44^, a U-Net architecture originally developed for non-human primate anatomical brain extraction. The model uses three adjacent slices as input and processes 256x256 rescaled patches. Data augmentation included random scaling, translation, flips, rotation, Gaussian noise, brightness and contrast jitter, and synthetic bias fields. The loss function combined Dice and cross-entropy losses with equal weights, and optimization used Adam with cosine annealing following warm-up epochs. The model achieved a validation Dice score of 0.95.

### Functional preprocessing

The functional workflow preprocesses BOLD and MION-based T2*-weighted time series.

The workflow includes slice timing correction when metadata are available, motion correction, within-session co-registration, bias correction of the temporal mean, conformation, brain extraction, and registration to T1w or template space.

Slice timing correction is performed with AFNI 3dTshift when BIDS metadata specify slice timing information. Motion correction is performed with FSL mcflirt using 6 DOF. The temporal mean of the motion-corrected time series serves as the representative functional image for subsequent steps. N4 bias field correction is applied to this representative image only, improving the reliability of downstream processing steps. The representative functional image is conformed using the same general strategy as the structural workflow, producing the FUNC reference image and defining FUNC space in the pipeline. A functional brain extraction CNN is subsequently applied to generate a brain mask, which is then applied to the 4D functional data.

Functional registration depends on the availability of a subject-specific T1w image. When a T1w image is available, the functional reference is registered to T1w space and then projected to template space by reusing the T1w-to-template transforms generated during the structural workflow. When no subject-specific T1w image is available, the functional reference is registered directly to the selected template.

### Functional validation analyses

Task-evoked validation was performed on the visual localizer dataset that used a block-design with four stimulus categories: scenes, faces, bodies, and objects. Preprocessed functional time series were analyzed using a first-level general linear model (GLM) implemented in Nilearn^45^. Each stimulus category was modeled using a canonical MION hemodynamic response function (HRF) parameterized as a gamma distribution, along with its time derivative. Nuisance regressors included cosine-basis drift terms (high-pass cutoff: 0.01 Hz), and temporal autocorrelation was modeled with an AR(1) noise model. The GLM was fit jointly across runs. Category-selective responses were estimated by contrasting each stimulus category against all remaining categories. Contrast maps were projected to the cortical surface and displayed at the top 5% of vertices for Fig. 4B.

Resting-state validation was performed on preprocessed resting-state data from the datasets described above. Regional time series were extracted from ARM2 parcels after registration to template space. For each monkey, Pearson correlations were computed between all pairs of regional time series to generate a region-by-region correlation matrix. Group-level correlation structure was summarized by averaging matrices across monkeys. Seed-based correlation maps were generated by correlating the time series from selected seed regions with all other cortical locations. To quantify similarity of resting-state organization across monkeys, each monkey’s regional correlation matrix was vectorized, and pairwise similarity was computed as the Pearson correlation between vectorized matrices. Similarity values were grouped according to whether monkey pairs came from the same dataset or from different datasets.

### Toolchain for main preprocessing steps

Brainana is implemented in Python and interfaces with established neuroimaging software packages for volumetric and surface preprocessing. Volumetric steps call FSL ^46^, AFNI ^47,48^, ANTs ^49^, and FireANTs ^50^; surface reconstruction calls FreeSurfer ^39^ and FastSurfer ^15^. The full toolchain is summarized in Table 1.

**Table 1.**
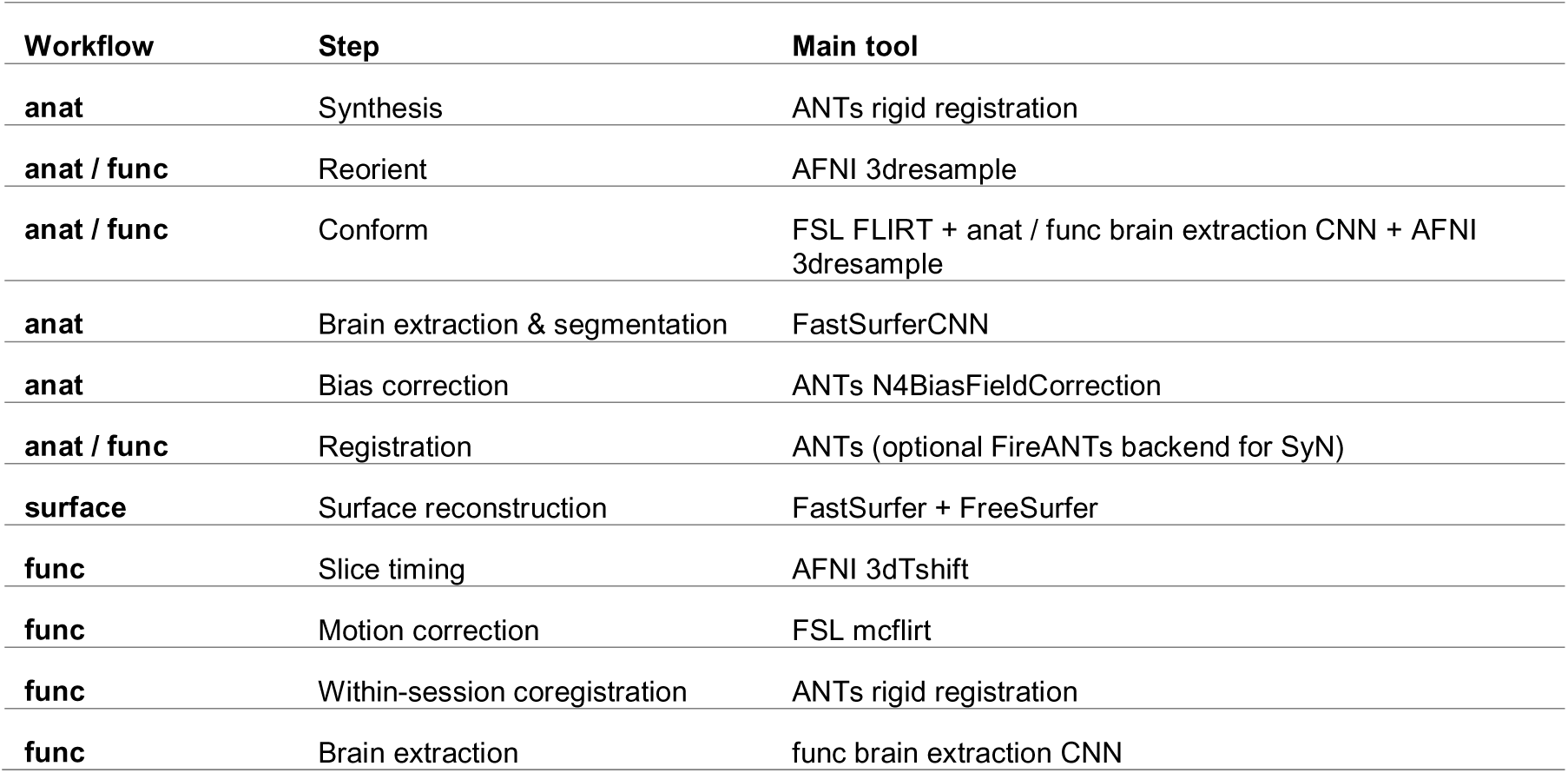
Brainana preprocessing toolchain. The table lists the major anatomical, functional, and surface reconstruction steps in Brainana and the primary software tools used to implement each step.

| Workflow | Step | Main tool |
| --- | --- | --- |
| anat | Synthesis | ANTs rigid registration |
| anat / func | Reorient | AFNI 3dresample |
| anat / func | Conform | FSL FLIRT + anat / func brain extraction CNN + AFNI 3dresample |
| anat | Brain extraction & segmentation | FastSurferCNN |
| anat | Bias correction | ANTs N4BiasFieldCorrection |
| anat / func | Registration | ANTs (optional FireANTs backend for SyN) |
| surface | Surface reconstruction | FastSurfer + FreeSurfer |
| func | Slice timing | AFNI 3dTshift |
| func | Motion correction | FSL mcflirt |
| func | Within-session coregistration | ANTs rigid registration |
| func | Brain extraction | func brain extraction CNN |

### Output structure

Brainana produces three categories of output: preprocessed derivatives, visual QC reports, and performance logs.

#### Preprocessed data

Outputs follow a BIDS derivatives layout. Structural outputs include preprocessed T1w and T2w images, brain-extracted images, brain masks, hemisphere masks, ARM2 segmentation, template-space images, atlas backprojections, and spatial transforms.

Functional outputs include preprocessed functional time series in scanner space, T1w space, and template space; brain masks; motion parameters; confound time series; and spatial transforms.

Surface outputs are written in FreeSurfer-compatible format and include cortical surface meshes, parcellations, and morphometric files such as thickness, area, and curvature. Example output structure is shown in Supp. Fig. 4.

#### Visual QC reports

For each subject, Brainana generates a browsable HTML report containing data summaries, QC snapshots, and a description of the preprocessing steps applied. These reports cover all major processing steps and are designed to facilitate rapid visual inspection of data quality. An example QC report is shown in Supp. Fig. 5.

#### Performance logs

Nextflow generates trace, timeline, and report files for each step, recording task status, exit code, submission time, wall-clock runtime, and resource usage. These logs support troubleshooting and resource planning across runs and sessions.

### Workflow management and containerization

To promote reproducibility, accessibility, and generalizability, Brainana uses Nextflow for workflow orchestration and Docker for containerized deployment. Nextflow manages dataset-scale execution by coordinating all preprocessing steps as a directed acyclic graph (DAG) of dependent processes, with individual steps implemented in Python or as calls to external tools. Nextflow handles input/output wiring, job scheduling, failure recovery, and resource allocation, supporting parallel execution across samples, transparent caching of completed steps, and seamless resumption of interrupted runs.

The pipeline is distributed as a Docker container that encapsulates all software dependencies at fixed versions. Containerization eliminates the need for users to install and configure individual dependencies and ensures a consistent runtime environment across systems, including laptops, and institutional clusters. It also supports reproducibility by allowing analyses to be rerun with the same software stack and pipeline version.

For users who require only volumetric T1w preprocessing of a single subject, or who lack access to a Docker-compatible environment, we provide Brainana Lite: a self-contained Jupyter notebook that runs the core structural pipeline. Brainana Lite is compatible with both local Jupyter and Google Colab, supports GPU acceleration, and requires only a T1w NIfTI file as input. It is intended for rapid prototyping and lightweight use cases; users requiring surface reconstruction, functional preprocessing, or full quality control should use the complete Brainana Docker pipeline.

### Brainana Viewer

Brainana Viewer is built on NiiVue^51^ and run locally through a WebGL2 compatible interface. The viewer interfaces directly with a Brainana output directory, whether stored locally or on a remote workstation accessed over SSH/SFTP, and automatically detects and loads each subject’s anatomical volumes, cortical surfaces, morphometric measures, and projected atlases, without requiring users to locate or configure individual derivative files. Multiplanar volume and 3D surface views are rendered as linked instances with shared crosshair coordinates, such that a location selected in one representation is reflected in the other. Categorical and continuous atlases, morphology overlays (curvature, sulcal depth, thickness), and functional maps (for example, retinotopic eccentricity and polar angle, with F-statistic thresholding) can be displayed on both volumes and surfaces, together with per-crosshair reports of coordinates, atlas labels, and functional values, and a visual-field plot for backprojected retinotopic maps. Brainana Viewer is distributed as a native desktop application for macOS, Linux, and Windows.

## Data Availability

The PRIME-DE and UNC-Wisconsin datasets are publicly available and can be individually accessed at the discretion of each originating study. The in-house dataset is available upon reasonable request to the corresponding author.

## Code Availability

Full documentation for Brainana is available at https://brainana.readthedocs.io/. The source code is available at https://github.com/xingyu-liu/brainana. The Docker container is available at https://hub.docker.com/repository/docker/liuxingyu987/brainana. Brainana Viewer is available at https://github.com/arcaro-lab/brainana_tools. Templates, atlases, and map zoo files are available at https://github.com/xingyu-liu/brainana/tree/main/template_zoo. Template surfaces are available at https://github.com/xingyu-liu/macaque_template_surfaces.

## Reference

1. Messinger, A. et al. A collaborative resource platform for non-human primate neuroimaging. NeuroImage 226, 117519 (2021).

2. Milham, M. P. et al. An Open Resource for Non-human Primate Imaging. Neuron 100, 61–74.e2 (2018).

3. Li, X. et al. Moving beyond processing- and analysis-related variation in resting-state functional brain imaging. *Nat*. Hum. Behav. 8, 2003–2017 (2024).

4. Taylor, P. A., et al. Editorial: Demonstrating quality control (QC) procedures in fMRI. Front. Neurosci. 17, 1205928 (2023)

5. Esteban, O. et al. fMRIPrep: a robust preprocessing pipeline for functional MRI. Nat. Methods 16, 111–116 (2019).

6. Ren, J. et al. DeepPrep: an accelerated, scalable and robust pipeline for neuroimaging preprocessing empowered by deep learning. Nat. Methods 22, 473–476 (2025).

7. Autio, J. A. et al. Towards HCP-Style macaque connectomes: 24-Channel 3T multi-array coil, MRI sequences and preprocessing. NeuroImage 215, 116800 (2020).

8. Garcia-Saldivar, P. et al. PREEMACS: Pipeline for preprocessing and extraction of the macaque brain surface. NeuroImage 227, 117671 (2021).

9. Jung, B. et al. A comprehensive macaque fMRI pipeline and hierarchical atlas. Neuroimage 235, 117997 (2021).

10. Kindred, N. et al. AutoMacq: an automatic pipeline to analyse macaque structural MRI data. bioRxiv 2023.06. 06.543846 (2023).

11. Lepage, C. et al. CIVET-Macaque: An automated pipeline for MRI-based cortical surface generation and cortical thickness in macaques. NeuroImage 227, 117622 (2021).

12. Tasserie, J. et al. Pypreclin: An automatic pipeline for macaque functional MRI preprocessing. NeuroImage 207, 116353 (2020).

13. Gorgolewski, K. J. et al. BIDS apps: Improving ease of use, accessibility, and reproducibility of neuroimaging data analysis methods. PLoS Comput. Biol. 13, e1005209 (2017).

14. Di Tommaso, P. et al. Nextflow enables reproducible computational workflows. Nat. Biotechnol. 35, 316–319 (2017).

15. Henschel, L. et al. Fastsurfer-a fast and accurate deep learning based neuroimaging pipeline. NeuroImage 219, 117012 (2020).

16. Hartig, R. et al. Subcortical Atlas of the Rhesus Macaque (SARM) for magnetic resonance imaging. bioRxiv 2020.09. 16.300053 (2020).

17. Dale, A. M., Fischl, B. & Sereno, M. I. Cortical surface-based analysis: I. Segmentation and surface reconstruction. Neuroimage 9, 179–194 (1999).

18. Alldritt, S. et al. Brain charts for the rhesus macaque lifespan. BioRxiv (2024).

19. Franklin, M. S. et al. Gender differences in brain volume and size of corpus callosum and amygdala of rhesus monkey measured from MRI images. Brain Res. 852, 263–267 (2000).

20. Tan, Z. et al. Brain development during the lifespan of cynomolgus monkeys. Neuroimage 305, 120952 (2025).

21. Hayashi, T. et al. The nonhuman primate neuroimaging and neuroanatomy project. Neuroimage 229, 117726 (2021).

22. Xia, J., Wang, F., Wang, Y., Wang, L. & Li, G. Longitudinal mapping of the development of cortical thickness and surface area in rhesus macaques during the first three years. Proc. Natl. Acad. Sci. 120, e2303313120 (2023).

23. Arcaro, M. J., Schade, P. F., Vincent, J. L., Ponce, C. R. & Livingstone, M. S. Seeing faces is necessary for face-domain formation. Nat. Neurosci. 20, 1404–1412 (2017).

24. Kornblith, S., Cheng, X., Ohayon, S. & Tsao, D. Y. A Network for Scene Processing in the Macaque Temporal Lobe. Neuron 79, 766–781 (2013).

25. Pinsk, M. A. et al. Neural Representations of Faces and Body Parts in Macaque and Human Cortex: A Comparative fMRI Study. J. Neurophysiol. 101, 2581–2600 (2009).

26. Popivanov, I. D., Jastorff, J., Vanduffel, W. & Vogels, R. Stimulus representations in body-selective regions of the macaque cortex assessed with event-related fMRI. NeuroImage 63, 723–741 (2012).

27. Tsao, D. Y., Freiwald, W. A., Knutsen, T. A., Mandeville, J. B. & Tootell, R. B. H. Faces and objects in macaque cerebral cortex. Nat. Neurosci. 6, 989–995 (2003).

28. Seidlitz, J. et al. A population MRI brain template and analysis tools for the macaque. NeuroImage 170, 121–131 (2018).

29. Balan, P. F. et al. MEBRAINS 1.0: A new population-based macaque atlas. Imaging Neurosci. 2, imag–2–00077 (2024).

30. Donahue, C. J. et al. Using Diffusion Tractography to Predict Cortical Connection Strength and Distance: A Quantitative Comparison with Tracers in the Monkey. J. Neurosci. 36, 6758–6770 (2016).

31. Saleem, K. S. et al. High-resolution mapping and digital atlas of subcortical regions in the macaque monkey based on matched MAP-MRI and histology. NeuroImage 245, 118759 (2021).

32. Lu, Y. et al. Macaque Brainnetome Atlas: A multifaceted brain map with parcellation, connection, and histology. Sci. Bull. 69, 2241–2259 (2024).

33. Thomas Yeo, B. T., et al. The organization of the human cerebral cortex estimated by intrinsic functional connectivity. J. Neurophysiol. 106, 1125–1165 (2011).

34. Xu, T. et al. Cross-species functional alignment reveals evolutionary hierarchy within the connectome. NeuroImage 223, 117346 (2020).

35. Arcaro, M. J. & Livingstone, M. S. Retinotopic Organization of Scene Areas in Macaque Inferior Temporal Cortex. J. Neurosci. 37, 7373–7389 (2017).

36. Arcaro, M. J., Schade, P. F. & Livingstone, M. S. Body map proto-organization in newborn macaques. Proc. Natl. Acad. Sci. 116, 24861–24871 (2019).

37. Oishi, H., Berezovskii, V. K., Livingstone, M. S., Weiner, K. S. & Arcaro, M. J. Metabolic organization of macaque visual cortex reflects retinotopic eccentricity and category selectivity. Preprint at 10.1101/2025.09.27.678945 (2025).

38. Arcaro, M. J., Ponce, C. & Livingstone, M. The neurons that mistook a hat for a face. eLife 9, e53798 (2020).

39. Fischl, B., Sereno, M. I., Tootell, R. B. H. & Dale, A. M. High-resolution intersubject averaging and a coordinate system for the cortical surface. Hum. Brain Mapp. 8, 272–284 (1999).

40. Feilong, M., Jiahui, G., Gobbini, M. I. & Haxby, J. V. A cortical surface template for human neuroscience. Nat. Methods 21, 1736–1742 (2024).

41. Ciric, R. et al. TemplateFlow: FAIR-sharing of multi-scale, multi-species brain models. Preprint at 10.1101/2021.02.10.430678 (2021).

42. Livingstone, M. S. et al. Development of the macaque face-patch system. Nat. Commun. 8, 14897 (2017).

43. Young, J. T. et al. The UNC-Wisconsin Rhesus Macaque Neurodevelopment Database: A Structural MRI and DTI Database of Early Postnatal Development. Front. Neurosci. 11, (2017).

44. Wang, X. et al. U-net model for brain extraction: Trained on humans for transfer to non-human primates. NeuroImage 235, 118001 (2021).

45. Nilearn contributors et al. nilearn. Zenodo 10.5281/ZENODO.8397156 (2026).

46. Jenkinson, M., Beckmann, C. F., Behrens, T. E. J., Woolrich, M. W. & Smith, S. M. FSL. NeuroImage 62, 782–790 (2012).

47. Cox, R. W. AFNI: Software for Analysis and Visualization of Functional Magnetic Resonance Neuroimages. Comput. Biomed. Res. 29, 162–173 (1996).

48. Cox, R. W. & Hyde, J. S. Software tools for analysis and visualization of fMRI data. NMR Biomed. 10, 171–178 (1997).

49. Avants, B., Epstein, C., Grossman, M. & Gee, J. Symmetric diffeomorphic image registration with cross-correlation: Evaluating automated labeling of elderly and neurodegenerative brain. Med. Image Anal. 12, 26–41 (2008).

50. Jena, R., Chaudhari, P. & Gee, J. C. FireANTs: Adaptive Riemannian Optimization for Multi-Scale Diffeomorphic Matching. Preprint at 10.48550/ARXIV.2404.01249 (2024).

51. Taylor Hanayik et al. niivue/niivue: @niivue/niivue-v0.69.0. Zenodo 10.5281/ZENODO.20310755 (2026).

