## Supplemental figures for "Brainana: an end-to-end preprocessing framework for macaque neuroimaging"

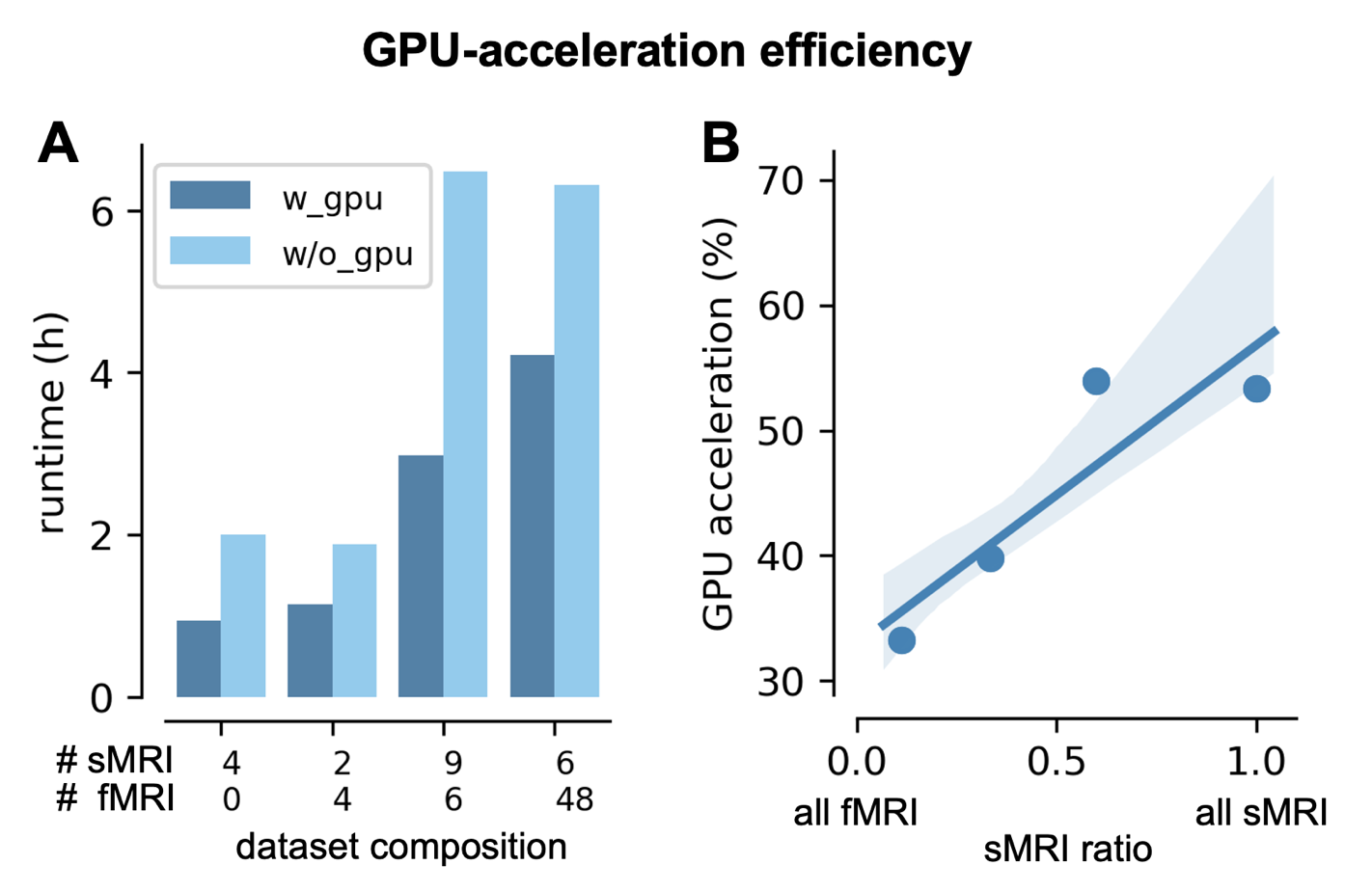

**Supp. Fig. 1 GPU-acceleration efficiency varies with dataset composition.**

**(A)** Runtime comparison with and without GPU acceleration across datasets containing different proportions of sMRI and fMRI data, ranging from sMRI only to fMRI only. Processing was benchmarked on a workstation with an AMD Threadripper 7995X CPU using 8 cores; RAM, 20 GB; GPU, NVIDIA RTX A6000 (48 GB VRAM).

**(B)** Linear regression of GPU-driven speedup (% runtime reduction) as a function of anatomical-to-functional image ratio (shaded region, 95% confidence interval). GPU acceleration scales positively with anatomical content, yielding approximately 60% runtime reduction for anatomical-only datasets and approximately 35% for functional-only datasets.

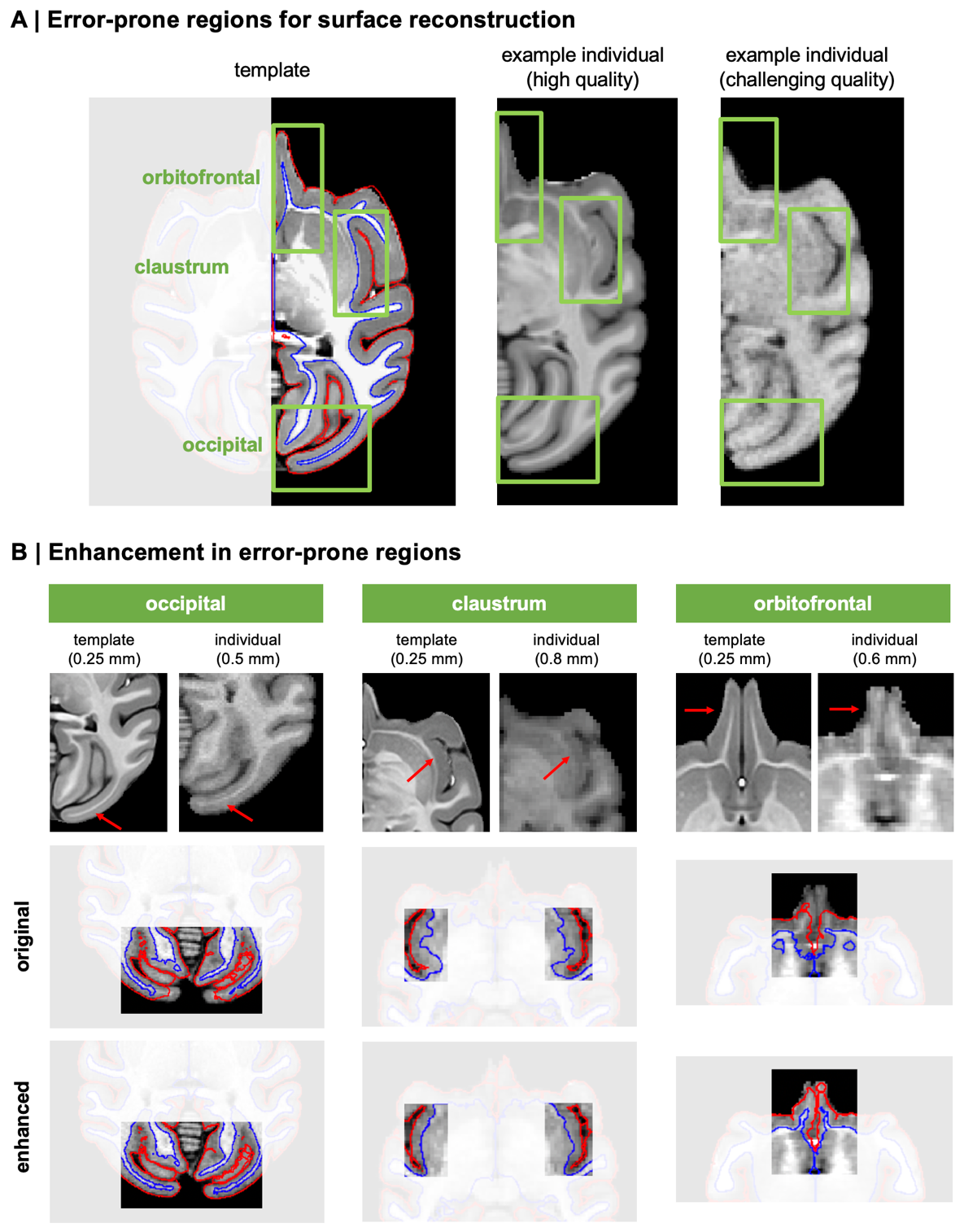
**Supp. Fig. 2. Targeted refinements improve surface reconstruction in challenging macaque cortical regions**

**(A) Cortical regions that commonly produce surface reconstruction errors in macaque MRI**. The NMT2Sym template is shown in sagittal view with reconstructed white matter (red) and pial (blue) surfaces overlaid. Green boxes highlight three regions that commonly require macaque-specific handling: orbitofrontal cortex (top), claustrum (middle), and the calcarine cortex (bottom). Corresponding zoomed views of each region are shown for a high-quality individual acquisition and a challenging-quality acquisition, illustrating the variability in image contrast and tissue boundary definition that drives reconstruction errors in these areas.

**(B) Effects of targeted surface reconstruction refinements**. For occipital, claustrum, and orbitofrontal regions, template and individual images are shown alongside surfaces before (original) and after (enhanced) the macaque-specific refinement. Red arrows indicate structures prone to segmentation of reconstruction errors. The enhanced workflow improves delineation of white matter and pial surfaces in regions where thin white matter, low tissue contrast, or nearby subcortical structures can disrupt reconstruction.

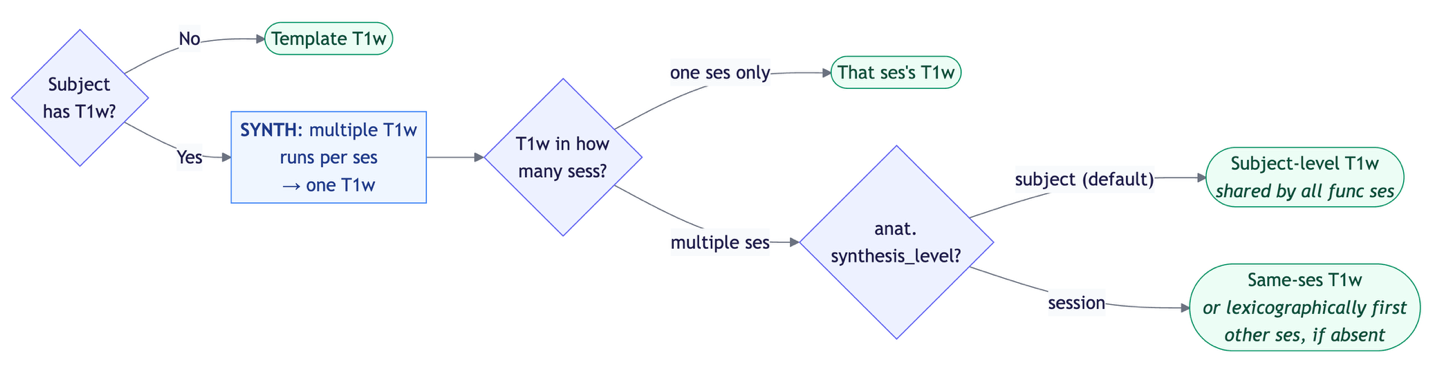

**Supp. Fig. 3 Decision logic for anatomical reference selection during functional processing.**

Flowchart depicting the decision logic by which Brainana assigns a structural reference for each functional session. When no subject-specific T1w image is available, functional data are registered directly to the selected template T1w image. When one or more subject-specific T1w images are available, multiple runs acquired within the same session are first synthesized into a single T1w reference. If T1w data span multiple sessions, the *anat.synthesis_level* configuration parameter governs alignment strategy: functional data are registered through a shared subject-level T1w reference by default, which is appropriate for cross-sectional designs, or through a session-specific T1w reference for longitudinal designs.

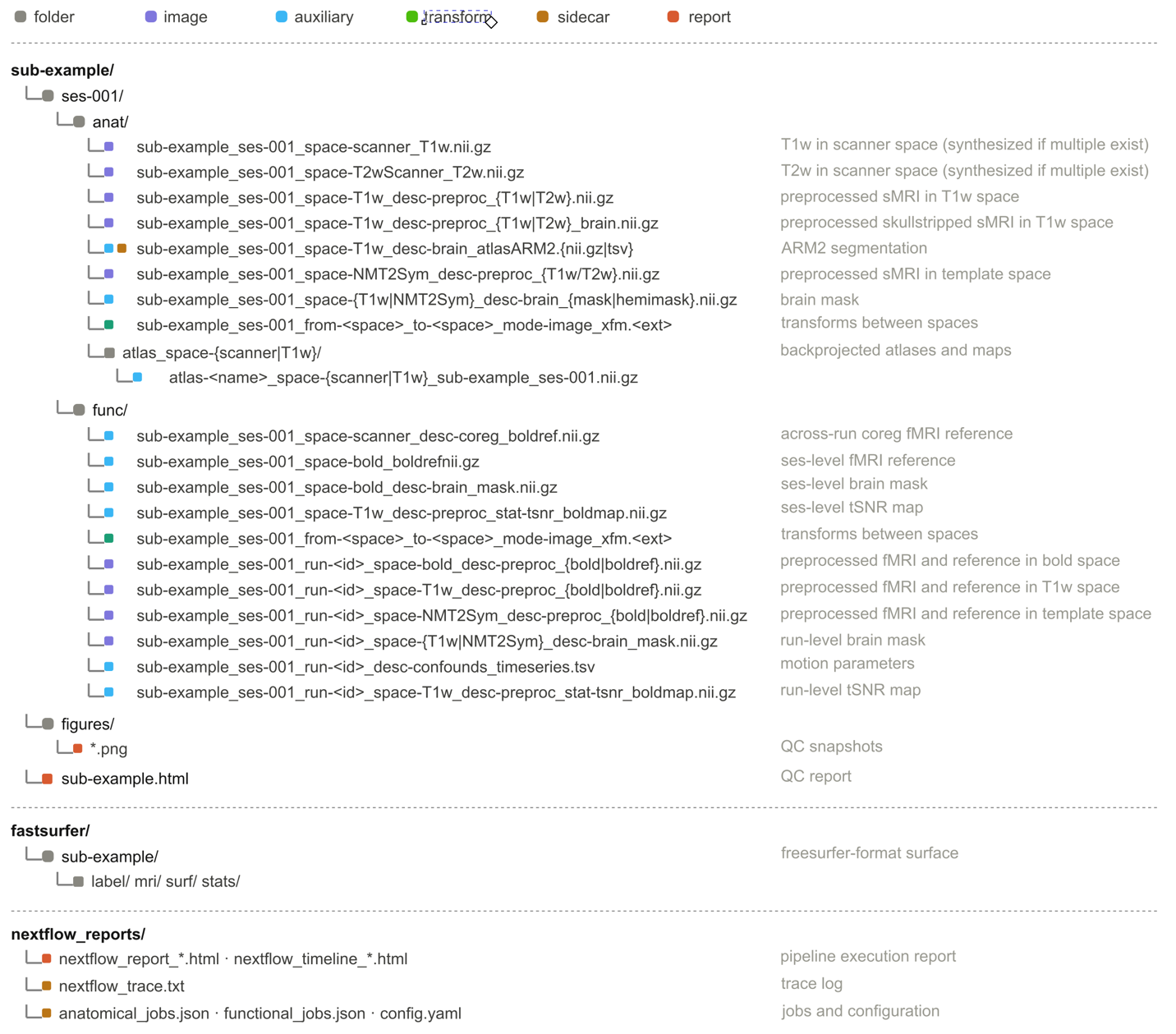

**Supp. Fig. 4. Canonical layout of Brainana output.**

Example Brainana derivatives layout showing the organization of preprocessed outputs, spatial transforms, surface files, quality-control reports, and performance logs. Structural derivatives include preprocessed T1w and optional T2w images in T1w and template spaces, brain masks, hemisphere masks, ARM2 segmentations, atlas back-projections, and inter-space transforms. Functional derivatives include preprocessed time series in scanner, T1w, and template spaces, brain masks, motion parameters, and inter-space transforms. Surface derivatives are written in FreeSurfer-compatible and GIFTI formats and include cortical meshes, parcellations, and morphometric measures such as cortical thickness, surface area, and curvature. Quality-control outputs include per-session PNG snapshots and an HTML report, and pipeline logs include Nextflow execution reports and job configuration files.

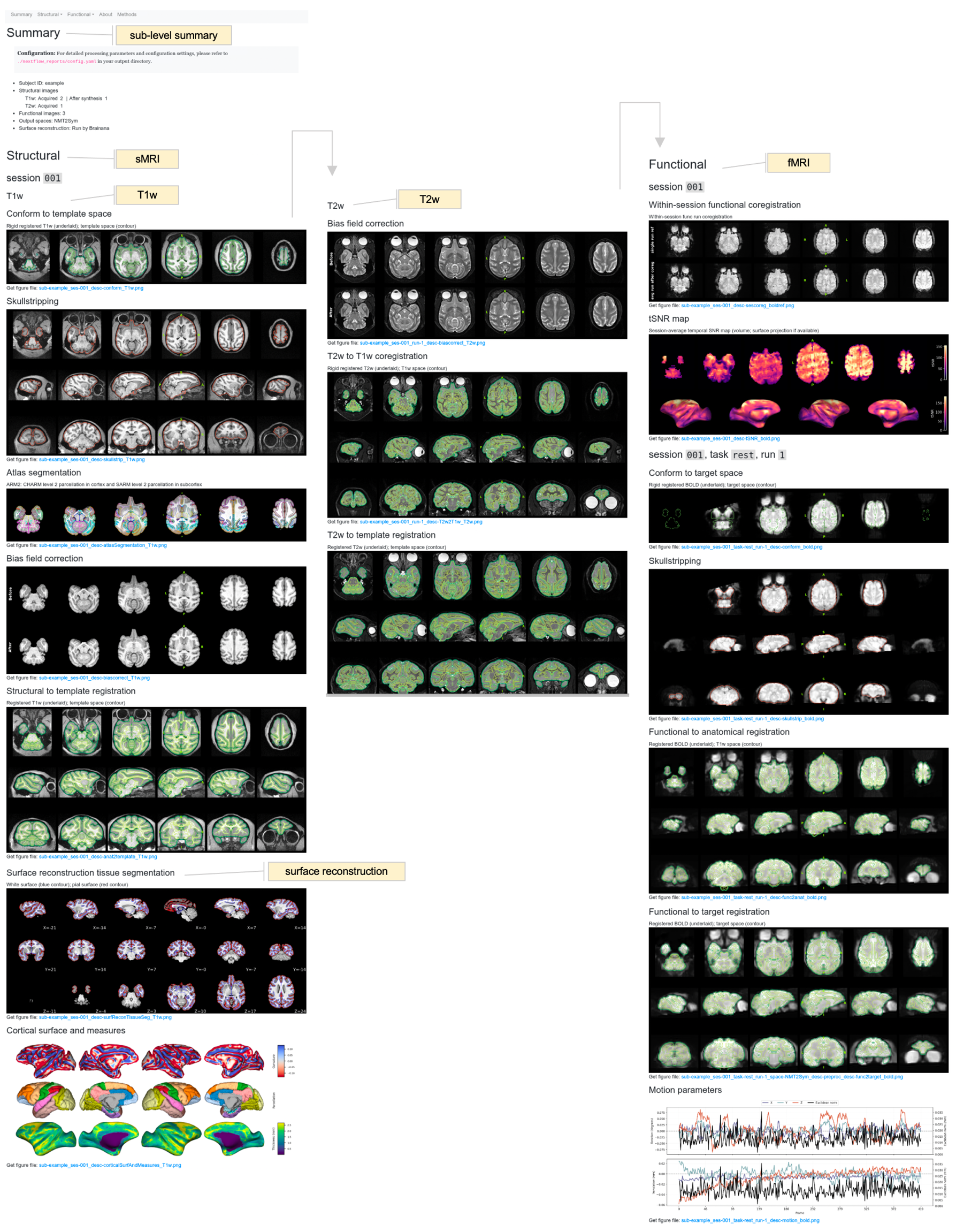

**Supp. Fig. 5 Canonical Brainana quality control report.**

Example HTML quality control report generated by Brainana for visual inspection of preprocessing outputs. Reports summarize the input data, processing steps, and key derivatives for each subject or session, and include snapshot overlays for major workflow stages such as structural conformation, brain extraction, tissue segmentation, registration, functional alignment, and surface reconstruction. These reports provide a standardized interface for evaluating preprocessing quality, identifying potential failures, and documenting processing outcomes across datasets.
